# Playing soft with cooperators emerges as a moral norm and promotes cooperation in evolutionary games

**DOI:** 10.1101/2021.03.01.433361

**Authors:** Mohammad Salahshour

## Abstract

In many biological populations, individuals face a complex strategic setting, where they need to make strategic decisions over a diverse set of issues. To study evolution in such a complex strategic context, here we introduce evolutionary models where individuals play two games with different structures. Individuals decide upon their strategy in a second game based on their knowledge of their opponent’s strategy in the first game. By considering a case where the first game is a social dilemma, we show that, as long as the second game has an asymmetric Nash equilibrium, the system possesses a spontaneous symmetry-breaking phase transition above which the symmetry between cooperation and defection breaks. A set of cooperation supporting moral norms emerges according to which cooperation stands out as a valuable trait. Notably, the moral system also brings a more efficient allocation of resources in the second game. This observation suggests a moral system has two different roles: Promotion of cooperation, which is against individuals’ self-interest but beneficial for the population, and promotion of organization and order, which is at both the population’s and the individual’s self-interest. Interestingly, the latter acts like a Trojan horse: Once established out of individuals’ self-interest, it brings the former with itself. Furthermore, we show that in structured populations, recognition noise can have a surprisingly positive effect on the evolution of moral norms and facilitates cooperation in the Snow Drift game.

## Introduction

Although beneficial for the group, cooperation is costly for the individuals, and thus, constitutes a social dilemma: Following their self-interest, individuals should refrain from cooperation. This leaves everybody worse off than if otherwise, all had cooperated [1–3]. Indirect reciprocity is suggested as a way out of this dilemma [3–6], which can also bring insights into the evolution of morality [5, 6]. Most models of indirect reciprocity consider a simple strategic setting where individuals face a social dilemma, commonly modeled as a Prisoner’s Dilemma. Individuals decide upon their strategy in the social dilemma based on the reputation of their opponent. In turn, reputation is built based on the strategy of the individuals in the same social dilemma. This self-referential structure can give rise to some problems. The core of these problems relies on how to define good and bad. In the simplest indirect reciprocity model, only first-order moral assessment rules are allowed: For instance, an individual’s reputation is increased by cooperation, and it is decreased by defection [7, 8]. This leads to a situation where defection with someone with a bad reputation leads to a bad reputation. From a moral perspective, this does not make sense. Besides, this can lead to the instability of the dynamic [5]. To solve these problems, it is possible to consider second-order moral assessment rules [9–13]. In prescribing an individual’s reputation, besides its action, second-order rules also take the reputation of its opponent into account. However, this way opens the door to third-order and higher-order moral assessment rules [14–17], which require having information about the actions of the individuals further and further into the past [5, 17, 18]. This leads to a rapid increase in the number and complexity of moral assessment rules by going to higher-order rules, even when, as it is commonly assumed, moral assessment is reduced to a binary world of good and bad [5, 15–19].

In contrast to the premise of most models of indirect reciprocity, strategic interactions in many biological contexts are not simple. Some kinds of strategic complexity have been considered in evolutionary games. Examples include strategy dependent stochastic transitions between social dilemmas [20, 21], deterministic transitions between games [22, 23], the dynamics of two or more evolutionary games played in parallel, in one shot [24–28] (also coined multigames [26]), or repeated interactions [29], heterogeneity in the payoff structure of the games played by the individuals [30–33], and the interaction of social dilemmas with signaling games [34–36]. Here we consider a strategically complex situation where individuals in a population face different strategic settings, each with a different set of actions and outcomes. In the language of game theory, this is to say individuals play different games, with different strategies and payoffs. In such a context, the individuals’ strategic decisions in one game can depend on what happens in the other games. This provides a way to solves the self-referential problem in the models of indirect reciprocity, as it allows the reputation-building mechanism and the decision-making mechanism to occur at different levels, i.e., in different games. As we show here, this observation can give rise to the evolution of a set of cooperation supporting moral norms.

In our model, individuals play a Prisoner’s Dilemma, together with a second game, which we call game *B*, and is not necessarily a social dilemma. Individuals build a reputation based on their behavior in the Prisoner’s Dilemma and act based on this reputation in the game *B*. We consider a situation where both games are played with the same opponent or when the two games are played with different opponents. When game *B* is a Prisoner’s Dilemma, or when it has a symmetric Nash equilibrium, the same problem incurred by indirect reciprocity models prevents the evolution of cooperation: Individuals are better off playing the Nash equilibrium, no matter what is the reputation of their opponent. The situation changes when game *B* has an asymmetric Nash equilibrium. In this case, it is in an individual’s best interest to take its opponent’s reputation into account in decision making. As we show here, this leads to the emergence of a symmetry-breaking phase transition above which the symmetry between cooperation and defection breaks. A set of behavioral rules emerges according to which cooperation stands out as a valuable trait, and individuals play softly with cooperators. This leads to the evolution of cooperation. Importantly, this set of rules also promote a more efficient allocation of resources in game *B*. This is particularly the reason why it evolves based on individuals’ self-interest. This observation appears to conform to the view that many aspects of moral systems do not necessarily require self-sacrifice but simply help to foster mutualistic cooperation and bring order and organization into societies [37–41]. In this regard, our analysis suggests that moral systems act as a Trojan horse: Once established out of the individual’s self-interest, they promote cooperation and self-sacrifice too.

Analysis of the model in a structured population shows that the results are robust in a structured population, as well. Furthermore, noise in inferring reputation can have a surprisingly positive effect on the evolution of a moral system in structured populations. However, this may not compensate the loss cooperators experience due to a high recognition noise level. Furthermore, we show a very high level of recognition noise facilitates the evolution of cooperative behavior in the snow-drift game. This contrasts previous findings regarding the detrimental effect of population structure on the evolution of cooperation in the snow-drift game in simple strategic settings [42], and parallels some arguments regarding the beneficial effect of noise for biological functions [43–46].

## The model

We begin by introducing two slightly different models. In the first model, the direct interaction Model, information about the strategies of the individuals is acquired through direct observation. In this model, we consider a population of *N* individuals. At each time step, individuals are randomly paired to interact. Each pair of individuals play a Prisoner’s Dilemma (PD) game, followed by a second game. The second game is a two-person, two-strategy game, which we call game *B*. We call the two possible strategies of game *B*, down (d), and up (u) strategies. The strategy of an individual in game *B* is a function of its opponent’s strategy in the first game. Thus, the strategy of an individual can be denoted by a sequence of three letters *abc*. Here, the first letter is the individual’s strategy in the PD and can be either *C* (cooperation) or *D* (defection). The second letter is the individual’s strategy in the game *B* if its opponent cooperates in the PD. The last letter is the individual’s strategy in the game *B* if its opponent defects in the PD. Clearly, we have *b, c* ∈ {*u, d*}. For example, a possible strategy is to cooperate in the PD, play *d* if the opponent cooperates, and play *u* if the opponent defects. We denote such a strategy by *Cdu*.

While in the first model individuals play both their games with the same opponent, we also consider a second model, the Reputation-based Model, where individuals play their two games with different opponents. In this model, at each time step, individuals are randomly paired to play a PD. After this, the interaction ends, and individuals meet another randomly chosen individual to play their second game. Under this scenario, individuals do not observe the strategy of their opponent in the PD. Instead, we assume individuals have a reputation of being cooperator or defector, on which, the decision of their opponent in the second game is based. For example, an individual with the strategy *Cdu*, cooperates in the PD, plays *d* if it perceives its opponent to be a cooperator, and plays *u* if it perceives its opponent to be a defector. To model reputation, we assume with probability 1 − *η* individuals guess the PD-strategy of their opponent correctly, and with probability *η* they make a mistake in guessing the PD-strategy of their opponent. *η* can be considered as a measure of noise in inferring the reputations. We note that, for *η* = 0, the dynamics of the two models are mathematically similar.

For the evolutionary dynamics, we assume individuals gather payoff according to the payoff structure of the games and reproduce with a probability proportional to their payoff. Offspring inherit the strategy of their parent. However, with probability *ν* a mutation occurs, in which case the strategy of the offspring is set to another randomly chosen strategy.

We will also consider a structured population. While in a mixed population individuals randomly meet to interact, in a structured population, individuals reside on a network and interact with their neighbors. That is, each individual derives payoffs by playing its two games with all its neighbors. For the evolutionary dynamics, we consider an imitation rule, in which individuals’ update their strategy in each evolutionary step by imitating the strategy of an individual in their extended neighborhood (composed of the individual and its neighbors) with a probability proportional to its payoff, subject to mutations. For the population network, we consider a first nearest neighbor square lattice with von Neumann connectivity and periodic boundaries.

For the PD, individuals can either cooperate or defect. If both cooperate, both get a payoff *R* (reward), and if both defect, both get a payoff *P* (punishment). If an individual cooperates while its opponent defects, the cooperator gets a payoff *S* (sucker’s payoff), while its opponent gets a payoff *T* (temptation). For a Prisoner’s Dilemma we have *S < P < R < T* with *T <* 2*R*. For game *B*, we show the payoff of mutual down by *R*_*B*_, and the payoff of mutual up by *P*_*B*_. If an individual plays up while its opponent plays down, the up-player gets *T*_*B*_ and the down-player gets *S*_*B*_.

We will analyze our model for different structures for game *B*. We begin by considering the cases where game *B* is a Snow Drift (SD) game (also known as the Hawk-Dove or Chicken game) [2], the Battle of the Sexes (BS), and the Leader game, and continue to examine the dependence of our results on the continuous variation of the structure of the game *B*. It is argued that these games, together with the Prisoners’ Dilemma constitute four archetypal two-person, two-strategy games, and encompass all the non-trivial distinct strategic situations [47]. We note that all these games have an asymmetric Nash equilibrium. In the Nash equilibrium, one of the strategies is superior in the sense that it leads to a higher payoff, leaving the opponent with a lower payoff. We call such a superior strategy the hard strategy, and the inferior strategy, which leads to a lower payoff in equilibrium, is called the soft strategy. When using the three archetypal games, we take the soft strategy to be the same as down, and the hard strategy to be the same as up.

## Results

### Mixed population

As shown in the Methods Section, the dynamics of the models can be described in terms of the replicator-mutator equations. We begin by setting *S* = 0, *P* = 1, and *R* = 3, and plot the density of cooperators in Fig. 1(a), and the density of the soft strategies in Fig. 1(b), as a function of *T*. Here from top to bottom, the second game is the SD, the BS, and the Leader game. See Table. 1 for the payoff values of game *B*. In each panel, we present the result of simulations (marker), together with the numerical solutions of the replicator dynamics (lines). The replicator dynamics show the model is bistable: Depending on the initial conditions, two fixed points, each with a high or a low level of cooperation are possible. These two fixed points are plotted by solid and dashed lines. The solid line represents the stationary state of the dynamics, starting from a random (unbiased) initial condition in which the density of all the strategies are equal. This can be considered as the equilibrium fixed point [48]. The dashed line represents the non-equilibrium fixed point, which can occur for certain initial conditions.

**Figure 1:**
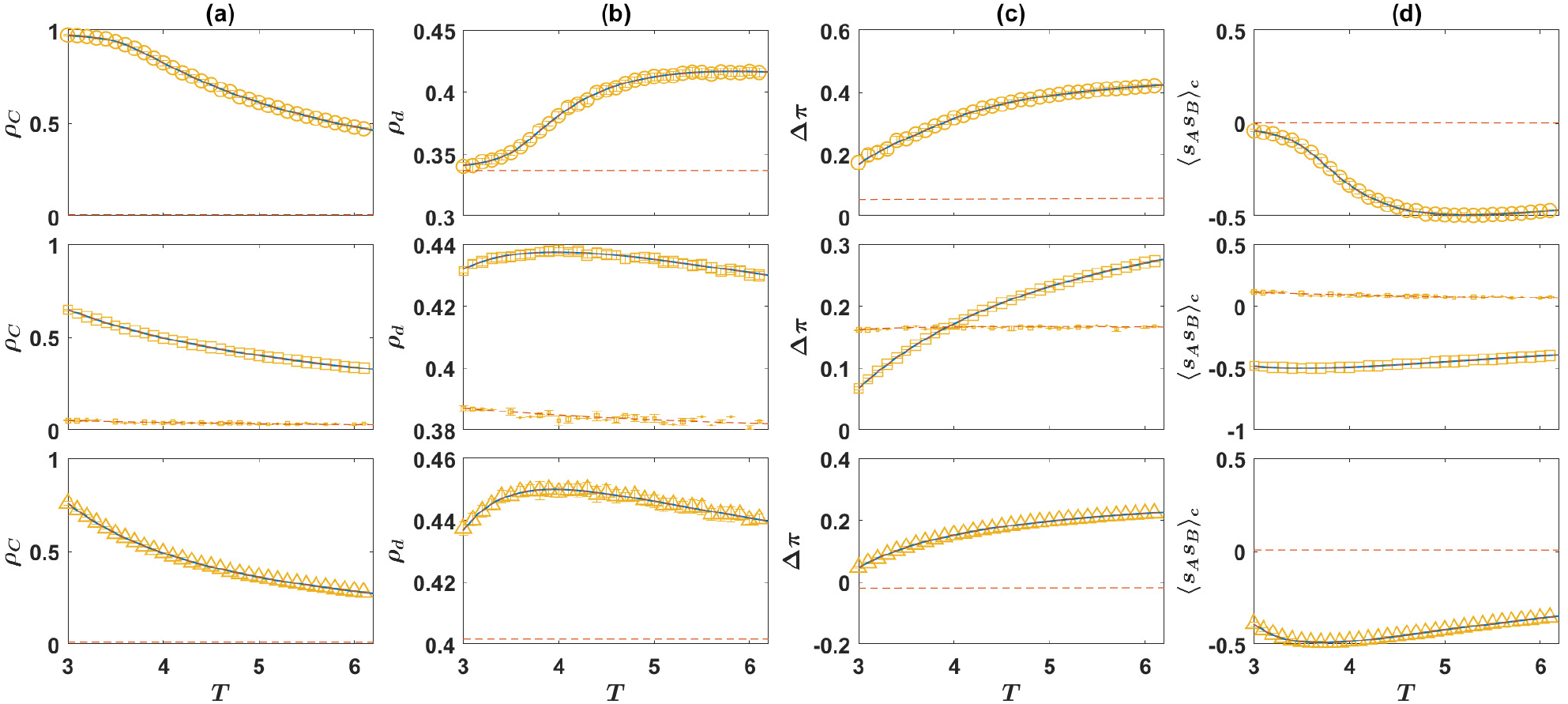
The direct interaction model with the three archetypal games. The density of cooperators (a), the density of soft strategies in game *B* (b), the normalized payoff difference of cooperators and defectors in game *B* (c), and the correlation between the strategy of the individuals in the two games (d), as a function of the temptation, *T*. Here, from top to bottom, the game *B* is the Snow Drift, the Battle of the Sexes, and the Leader game. The payoff values used for the games are presented in Table. 1. The lines show the result of the replicator dynamics, and the markers show the results of simulations. The solid blue line shows the equilibrium fixed point, which occurs starting from an unbiased initial condition in which the density of all the strategies are equal, and the dashed red line shows the non-equilibrium fixed point, which can occur starting from certain initial conditions. For the simulations, a sample of 80 simulations, in a population of size *N* = 10000 is used. The simulations start from random initial conditions. In each simulation, the dynamics settle in one of the two fixed points. The markers show the averages, and the error bars show the standard deviation in the sample of simulations that settle in the given fixed point, and the size of markers is proportional to the number of times that the given fixed point occurs. Here, *ν* = 0.005. The simulations are run for 20000 time steps, and an average over the last 1000 time steps is taken.

**Table 1:** Base payoff values.

| Prisoner's Dilemma |  |  |  |  | Game B |  |  |  |  |
| --- | --- | --- | --- | --- | --- | --- | --- | --- | --- |
| | $R$ | $S$ | $T$ | $P$ | | $R_B$ | $S_B$ | $T_B$ | $P_B$ |
| Prisoner's Dilemma | 3 | 0 | 5 | 1 | Snow Drift (SD) | 3 | 1 | 5 | 0 |
|  |  |  |  |  | Battle of the Sexes (BS) | 0 | 3 | 5 | 0 |
|  |  |  |  |  | Leader | 2 | 3 | 5 | 1 |

The markers represent the result of simulations in a population of size *N* = 10000. A set of 80 simulations is performed for *T* = 20000 time steps starting from random initial conditions (i.e. randomly assigning the strategies). In agreement with the results of the replicator dynamics, in each simulation, the dynamics settle in one of the two fixed points. The markers represent the mean over the realizations that settle in a given fixed point. The marker sizes are proportional to the number of times that a given stationary state has occurred in the set of 80 simulations.

As can be seen, the replicator dynamics predict starting from a random initial condition, for all the values of *T* the system settles in the cooperative fixed point. While this is often the case for a simulation in a finite population, depending on the structure of game *B*, with a low probability the non-equilibrium state, where a low level of cooperation is observed, can occur. This is the case for the BS games. The density of soft strategies, plotted in Fig. 1(b), is higher in the cooperative fixed point as well. On the other hand, in the non-cooperative fixed point, the density of soft strategies is smaller and close to the Nash equilibrium of the corresponding games. This shows, the coupling between games, not only promotes cooperation in the PD, but also increases cooperative behavior in game *B*.

So far we have seen that when a social dilemma is coupled with a second game, cooperation evolves in both games. The coupling of the two games is brought about by the fact that in deciding about their strategy in the second game, individuals use information about the strategy of their opponent in the first game. As cooperators receive a lower benefit from the social dilemma, they can survive only if their payoff from game B compensates for the cost of cooperation they pay in the social dilemma. To see this is indeed the case, in Fig. 1(c), we plot the normalized payoff difference of cooperators and defectors in the second game,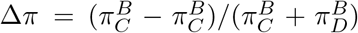. In all the three games, in the cooperative fixed point, cooperators receive a higher payoff in game *B*. In the SD and the BS games, cooperators receive a higher payoff in game *B*, even in the non-cooperative fixed point. The fact that cooperators’ payoff in game *B* is higher than that of the defectors indicates that in the course of evolution individuals develop strategies that tend to play soft with cooperators and hard with defectors. This can be considered as the emergence of a set of moral rules that allows cooperators to reach a benefit by being treated softly. Consequently, cooperators can reach a higher payoff by playing a hard strategy. Besides, by increasing the strength of the social dilemma (that is by increasing the temptation *T*), the payoff difference in game *B* increases in favor of the cooperators. This is due to an increase in the likelihood that cooperators receive a soft encounter in game *B*, by increasing the strength of the social dilemma (See the Supplementary Material, Fig. S.7). This shows the stronger a dilemma, and the higher the cost of cooperation, a stronger set of cooperation supporting moral norms emerge.

Emergence of a set of cooperation supporting moral norms can also lead to an anti-correlation between the strategies of the individuals in the two rounds: Cooperators are more likely to play a hard strategy in the second game, and defectors are more likely to play a soft strategy in the second game. This can be seen in Fig. 1(d), where the connected correlation of the strategies of the individuals in the two games, ⟨*s*_*A*_*s*_*B*_⟩_*c*_ = ⟨*s*_*A*_*s*_*B*_⟩ − ⟨*s*_*A*_⟩⟨*s*_*B*_⟩, is plotted. Here, ⟨·⟩ denotes an average over the population. To calculate the correlation function, we have assigned a value +1 to cooperation and the soft strategy, and −1 to the defection and hard strategies.

So far we have considered the direct interaction model, where individuals play both their games with the same opponent. The same phenomenon is at work, in the reputation-based model, where individuals play their two games with different opponents. In Fig. 2, we turn to the reputation-based model. In Fig. 2(a), we plot the density of cooperators in the Prisoner’s Dilemma, and in Fig. 2(b), we plot the density of the soft strategy in game *B*, as a function of the probability of error in inferring the PD-strategy of the opponent, *η*. Here, as before, from top to bottom, game *B* is, respectively, SD, BS, and the Leader game. Lines represent the results of the replicator dynamics, and markers show the results of simulations in a finite population. For a small probability of error, *η*, the model is bistable. The equilibrium fixed point is plotted by a solid blue line, and the non-equilibrium fixed point resulted from a biased initial condition, is plotted by dashed red line. However, for a large probability of error, the model becomes mono-stable and the dynamics settle in a defective fixed point in which the density of strategies in both games is close to their Nash equilibrium value. We note that, for *η* = 0.5, individuals have no information about the strategy of their opponent. As *η* increases beyond 0.5, individuals are more likely to infer the PD-strategy of their opponent erroneously than by chance. This gives net information that can be used by the population to self-organize in a cooperative fixed point. For this reason, the dynamic is symmetric around *η* = 0.5 and result in the same cooperation level for *η* and 1 − *η*.

**Figure 2:**
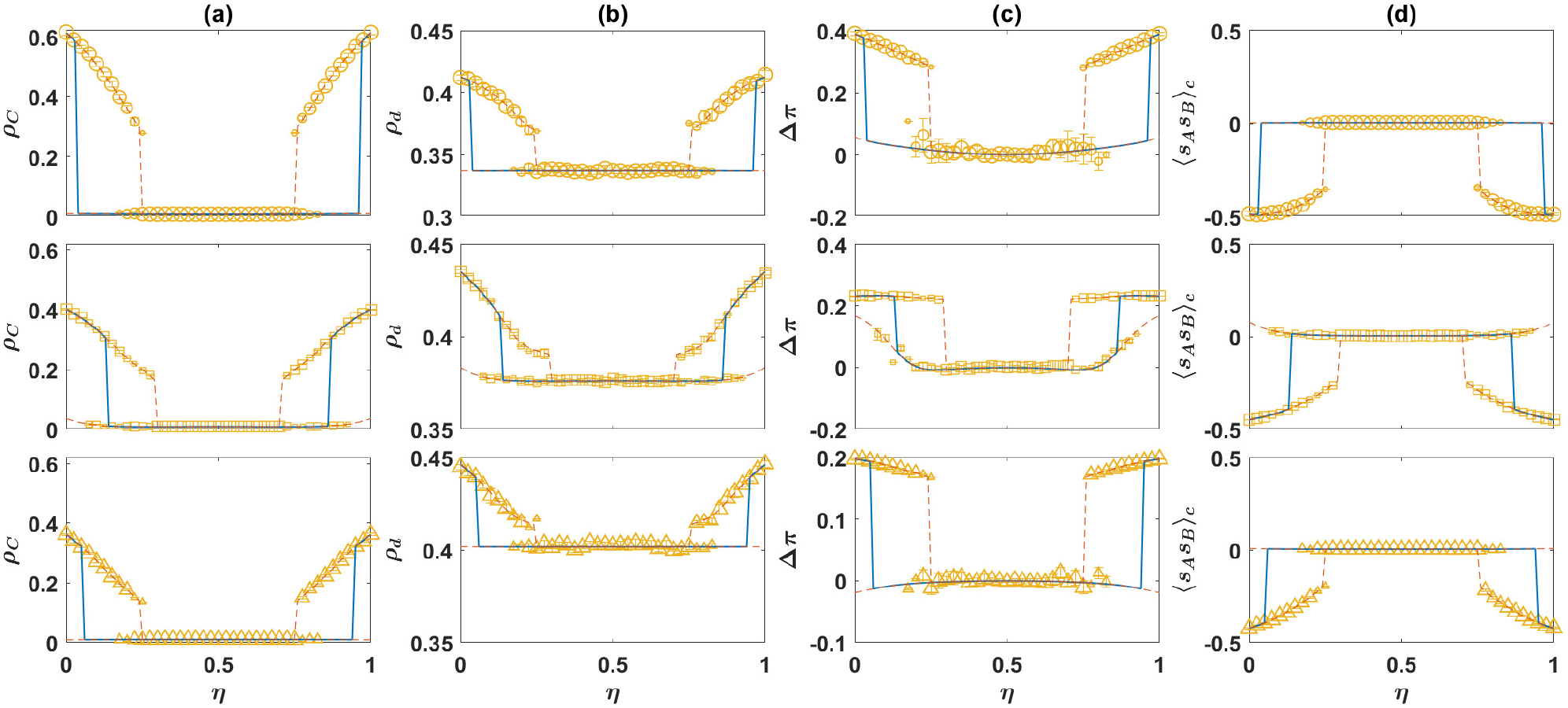
The reputation-based model with the three archetypal games. The density of cooperators (a), the density of soft strategies in game *B* (b), the normalized payoff difference of cooperators and defectors in game *B* (c), and the correlation between the strategy of the individuals in the two games (d), as a function of the probability of error in inferring the PD strategy of the opponent, *η*. Here, from top to bottom, the game *B* is the Snow Drift, the Battle of the Sexes, and the Leader game. The payoff values used for the games are presented in Table. 1. The lines show the result of the replicator dynamics, and the markers show the results of simulations. The solid blue line shows the equilibrium fixed point, which occurs starting from an unbiased initial condition in which the density of all the strategies are equal, and the dashed red line shows the non-equilibrium fixed point, which can occur for certain initial conditions. For the simulations, a sample of 80 simulations, in a population of size *N* = 10000 is used. The simulations start from random initial conditions. In each simulation, the dynamics settle in one of the two fixed points. The markers show the averages, and the error bars show the standard deviation in the sample of simulations that settle in the given fixed point. The size of the markers is proportional to the number of times that the given fixed point occurs. Here, *ν* = 0.005. The simulations are run for 20000 time steps, and an average over the last 1000 time steps is taken.

For the simulations, a sample of 80 simulations, run for 20000 time steps is used. Starting from a random initial condition, the dynamics settle in one of the stationary states. The size of the markers is proportional to the number of times that a given equilibrium occurs. As it is clear in the figure, finite size effects favor cooperation. This can be seen by noting that, in simulations in a finite population, the dynamics settle into the cooperative fixed point with a high probability, even when this is the non-equilibrium fixed point in the infinite size system.

The normalized payoff difference of cooperators and defectors in game 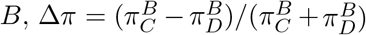, is plotted in Fig. 2(c). As can be seen, in the cooperative fixed point cooperators receive a higher payoff in game *B*. As in the direct interaction model, this is due to the emergence of a set of cooperation supporting moral norms, according to which, individuals are more likely to play the soft strategy with cooperators compared to defectors. This in turn allows cooperators, to be more likely to play a hard strategy in game *B*. This leads to an anti-correlation between the strategy of the individuals in the two games, as can be seen in Fig. 2(d). On the other hand, in the defective fixed point, the payoff difference of cooperators and defectors is near zero, and almost no correlation between the strategy of the individuals in the PD and game *B* is observed.

Thus far, using three different game structures for game *B*, we have seen when a social dilemma is accompanied by a second game, the dynamics can self-organize into a cooperative state where a set of cooperation supporting norms emerges and promotes cooperation. This raises the question that, are there certain conditions that the second strategic setting should satisfy for such norms to emerge? A characteristic of all these games is that they have an asymmetric Nash equilibrium. This is a necessary condition for the emergence of cooperation. When the second game has a symmetric equilibrium, individuals can maximize their payoff by consistently playing the Nash strategy in the second game, irrespective of the strategy of their opponent in the PD. This is particularly the reason why, when game *B* is a PD, cooperation in none of the games evolves. On the other hand, when the second game has an asymmetric Nash equilibrium, individuals need to coordinate on heterogeneous strategy pairs to maximize their own payoff. In this case, taking the strategy of their opponent in the first round into account can provide a way for them to efficiently play a strategy that increases their payoff.

To more clearly see how this happens, in Fig. 3, we plot the density of different strategies, in the direct interaction model, as a function of time. Here, game *B* is a Snow Drift game. The dynamic is similar for other games, and in the reputation-based model. Starting from a random initial condition, in which all the strategies are found in the same density, the population rapidly goes to a state where the density of strategies in both games is close to its Nash equilibrium value. This can be seen in Figs. 3(c) (replicator dynamics) and 3(d) (simulations in a population of size 10000), where the density of cooperators in the PD, and soft strategy in the SD are plotted. In this state, the densities of all the strategies are close to their value in the defective fixed point, such that it appears that the system is settled in the defective phase. However, this only sets the stage for the second phase of the evolution. As cooperators are found in a very small density in this phase, strategies that play softly with cooperators do not impose a large cost on their bearer and grow in number. When such strategies are accumulated enough, at a certain time the system shows a rapid dynamical transition to the cooperative fixed point where cooperators emerge in large numbers. Interestingly, cooperators always play a hard strategy with defectors, and defectors always play the soft strategy with cooperators. This compensates for the cost of cooperation that they pay. On the other hand, both cooperators and defectors play a combination of soft and hard strategies among themselves. This phenomenology shows, when a game has an asymmetric equilibrium, individuals can use information about the strategy of their opponent in a social dilemma, to efficiently coordinate in an asymmetric equilibrium and avoid paying the cost of coordination failure. Consequently, a set of behavioral or moral norms emerges, according to which cooperators are allowed to play hard, and deserve to be played soft with. This supports cooperation, by compensating for the cost of cooperation. Importantly, by facilitating coordination, this mechanism also increases the cooperation level in the second game.

**Figure 3:**
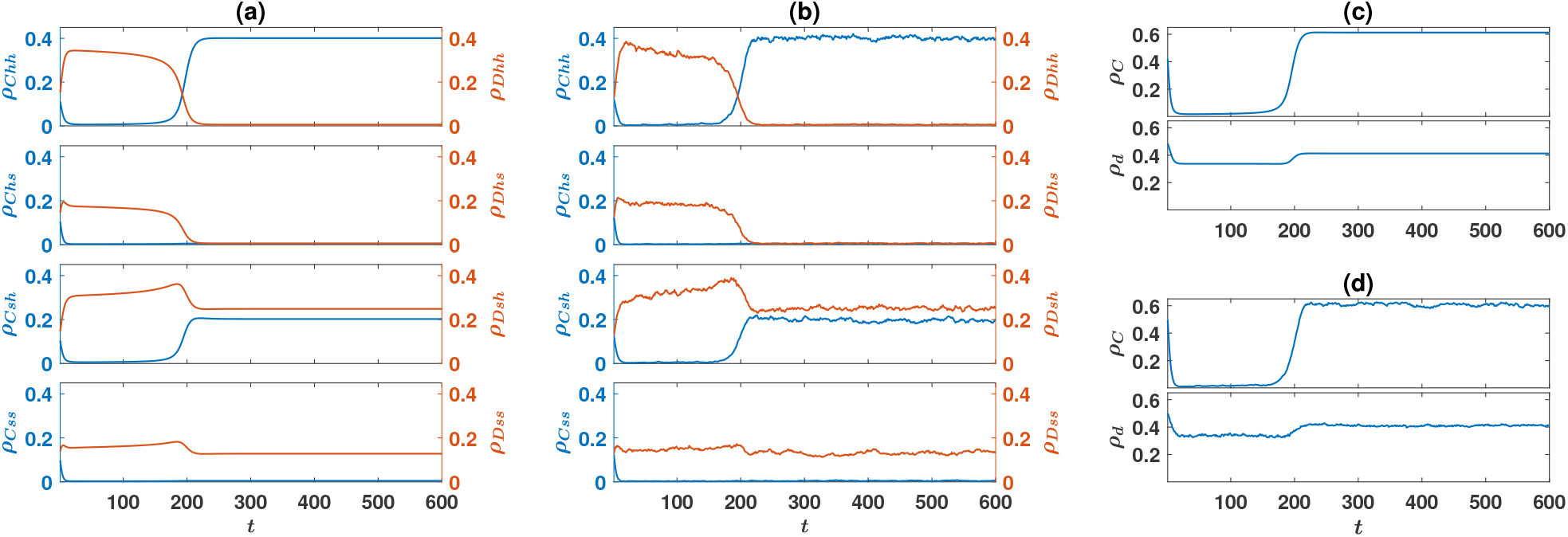
Time evolution in the model with direct interactions. (a) and (b): The time evolution of different strategies, resulted from the replicator dynamics (a), and a simulation in a finite population (b). (c) and (d): The time evolution of the density of the cooperators *ρ*_*C*_ (up), and the density of the soft strategies *ρ*_*d*_ (bottom), resulted from the replicator dynamics (c), and a simulation (d). The simulation is performed on a population of size *N* = 20000 and the mutation rate is *ν* = 0.005. The initial condition is a random assignment of strategies (for the replicator dynamics this implies *ρ*_*x*_ = 1*/*8, for all strategies *x*). The base payoff values presented in Table. 1, are used.

So far, using three archetypal games, we have seen cooperation and a set of cooperation supporting moral rules evolve in both models. To see how the models behave with respective to continuous variations of the structure of game *B*, we set *S* = 0, *P* = 1, *R* = 3, and *T* = 5 for the PD, and *R*_*B*_ = 3, and *P*_*B*_ = 1 for game *B*, and color plot the density of cooperators as a function of *S*_*B*_ and *T*_*B*_, in Fig. 4(a), for the direct interaction model, and in Fig. 4(b) for the reputation-based model. In the top panels, the results of the replicator dynamics are shown, and in the bottom panel, the results of simulations in a population of *N* = 1000 individuals are shown. Here a sample of *R* = 128 simulations, run for *T* = 10000 time steps is used. Averages are taken over the last 1000 time steps.

**Figure 4:**
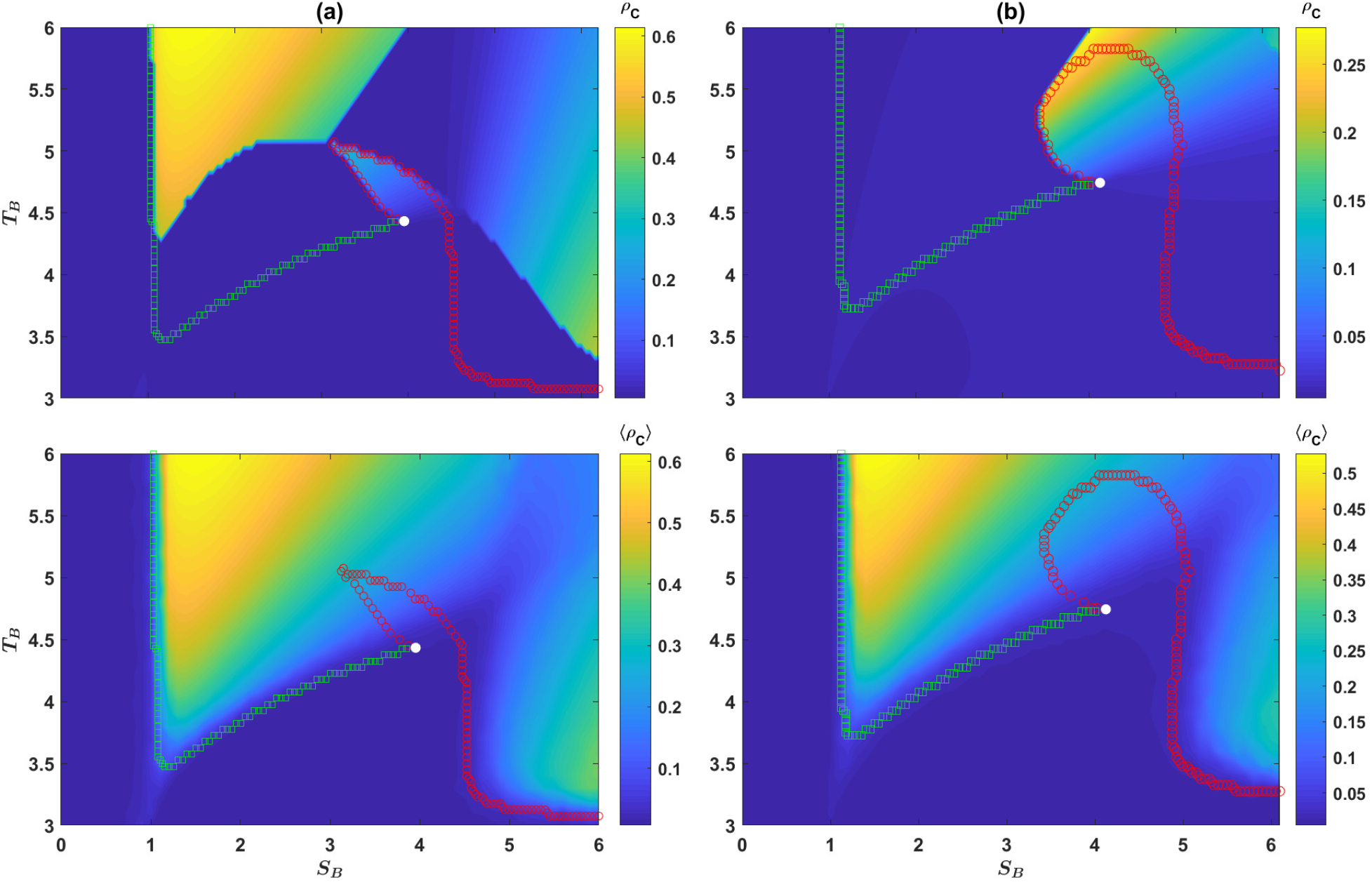
The behavior of the models under continuous variation of the structure of game *B*. The color plot of the density of cooperators, in the direct interaction model (a), and the reputation-based model (b), in the *S*_*B*_ − *T*_*B*_ plane. The top panels show the result of the replicator dynamics and the bottom panels show the results of simulations in a population of 1000 individuals. In both cases, an unbiased initial condition (random assignment of strategies) is used. We have set *R* = 3, *S* = 0, *P* = 1, *T* = 5, *R*_*B*_ = 3, and *P*_*B*_ = 1. The boundaries of bistability are plotted as well. Below this boundary the dynamic is monostable, settling into a fixed point with a low level of cooperation. Above the boundary, a cooperative fixed point becomes stable and the dynamics become bistable. The two branches of the boundary meet at a critical point, where the transition becomes continuous. A comparison shows finite size effects strongly favor cooperation. Here, *η* = 0.1, and *ν* = 0.005.

For *S*_*B*_ *<* 1, and *T*_*B*_ *> R*_*B*_ = 3, game *B* is a PD, with a symmetric Nash equilibrium. In this case, cooperation does not evolve. Similarly, for *T*_*B*_ *<* 3, game *B* has a symmetric Nash equilibrium. Cooperation does not evolve in this case either. On the other hand, for *T*_*B*_ *>* 3, and *S*_*B*_ *>* 1, game *B* has an asymmetric Nash equilibrium. In principle, cooperation can evolve in this case. The boundaries of bistability, plotted by markers, show the boundary above which the system becomes bistable. Below this boundary, only one fixed point, with a low level of cooperation is possible, and above this line, a cooperative fixed point emerges as well. In the bistable region, depending on the initial conditions, the dynamics settle in one of the two fixed points. The color plots show the cooperation level starting from unbiased initial conditions. A comparison of the results of the replicator dynamics and simulations in finite population shows finite size effects strongly favor cooperation, such that the transition between the two phases occurs for smaller values of *T*_*B*_ or *S*_*B*_ in a finite size population.

We note that the boundary of bistability is composed of two branches. Above each branch, a different fixed point occurs. Above the branch plotted by green squares, the soft strategy is *d* (for large *T*_*B*_), and above the branch plotted by red circles, the soft strategy is *u* (for large *S*_*B*_). The two branches meet at a single critical point, where the transition becomes a symmetry breaking continuous transition. Below the critical point, cooperation and defection in the PD are symmetric and are treated in the same way in game *B*. This leads to a symmetric state where the evolution of cooperation is prevented due to the cost of cooperation. Above the critical point, however, the symmetry between cooperative and defective strategies breaks, and a set of cooperation favoring norms emerges.

### Structured population

An interesting question is if strategic complexity can promote cooperation in structured populations too? It is well known that network reciprocity can promote cooperation under specific evolutionary processes [49, 50]. However, this is only the case when individuals are selected for reproduction with a probability proportional to the exponential of their payoff. In our model, individuals are selected for reproduction with a probability proportional to their payoff. In this case, network reciprocity can not promote cooperation when individuals play a Prisoner’s Dilemma. However, when the Prisoner’s Dilemma is followed by a second game, cooperation evolves in a structured population. This can be seen to be the case in Fig. 5(a) where the density of cooperators in the reputation-based model as a function of *η*, when game *B* is one of the three archetypal games, is plotted. This figure shows the results of simulations on a population of *N* = 40000 individuals residing on a 200×200 two-dimensional square lattice with first nearest neighbor von Neumann connectivity and periodic boundaries. Before proceeding to the analysis of the results, we note that, as explained before, *η* = 0.5 can be considered as the maximal noise level, where there is no net information in the inference of the individuals. Whereas, for *η* smaller or larger than 0.5, there is net information in the individuals’ inference, which can be used to increase payoffs. For this reason, in the following, we refer to the case of *η* = 0.5 as the maximal noise level.

**Figure 5:**
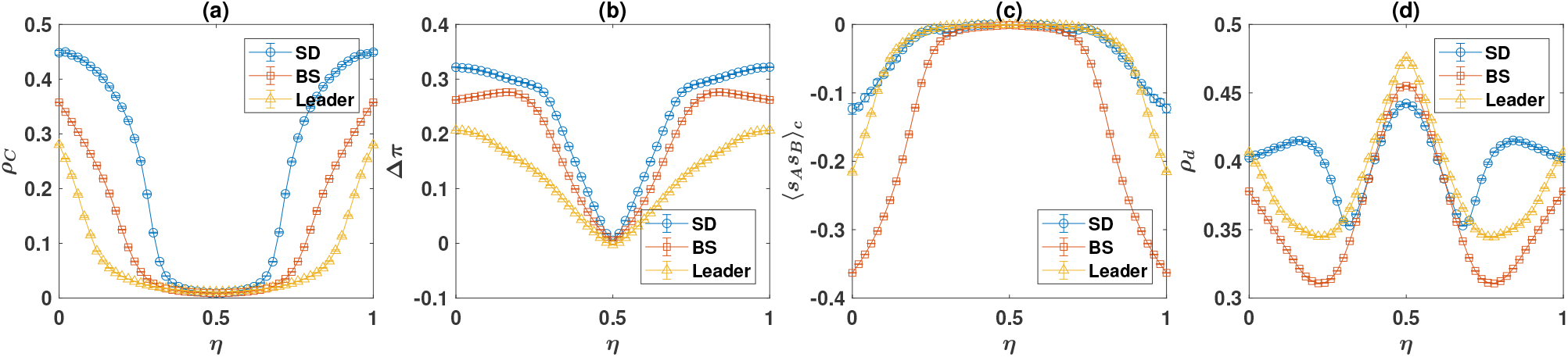
The reputation-based model with three archetypal games in a structured population. The density of cooperators (a), the normalized payoff difference of cooperators and defectors in game *B* (b), the correlation between the individuals’ strategies in the two games (c), and the density of soft strategies in game *B* (d), as a function of the probability of error in inferring the PD strategy of the opponent, *η*, are plotted. The payoff values used for the games are presented in Table. 1. Simulations are performed in a population of 40000 individuals residing on a 200 × 200 square lattice with first nearest neighbor von Neumann connectivity and periodic boundaries. The simulations are performed for 6000 time steps, and averages and standard deviations are calculated based on the last 4000 time steps. The simulations start from random initial conditions. Here, *ν* = 0.005.

Cooperation in the Prisoner’s Dilemma evolves as long as noise in recognition is small enough. As in the case of the mixed population, this is due to the evolution of cooperation supporting moral norms, which guarantee a higher payoff for cooperators in game *B*. This can be seen to be the case in Fig. 5(b), where the normalized payoff difference of cooperators and defectors in game *B* is plotted. Furthermore, due to the evolution of cooperation supporting norms, cooperators are more likely to play a hard strategy in game *B*, than the defectors are. This leads to the negativity of the connected correlation function of the individuals’ strategies in the two games, as shown in Fig.5(c).

The moral system also supports a high level of cooperative strategies in game *B*. This can be seen in Fig. 5(d), where the density of soft strategies in the population is plotted. The density of soft strategies is always larger than the Nash equilibrium value. This is particularly surprising, given that, although beneficial for the evolution of cooperation in the Prisoner’s Dilemma, network structure can hinder the evolution of cooperation in the Snow Drift game [42]. Our results show that in contrast to what is the case is a simple strategic setting, strategic complexity can provide an avenue for network structure to play a constructive role for the evolution of cooperative behavior in the Snow Drift game. Interestingly, the level of cooperative behavior in game *B* is maximized for maximal noise level. That is for *η* = 0.5, where there is no information in individuals’ recognition of others’ strategy in the Prisoner’s Dilemma.

To take a more in-depth look into the dynamics of the system, in Fig. 6, we plot the density of different strategies in the population, for the cases that the game *B* is one of the three archetypal games. As the probability of error approaches 0.5, cooperative strategies decrease in density. However, the density of defectors who in practice are more likely to defer to cooperators than not, that is *Ddu* for *η <* 0.5, and *Dud* for *η >* 0.5, increases when the noise level approaches the maximal value of 0.5. This observation shows that higher noise level can strengthen the moral system, such that cooperation supporting norms become stronger in higher noise level. This can be more clearly seen to be the case in Fig. 7, where the densities of the strategies who play up with cooperators, *u*(*C*), down with cooperators *d*(*C*), up with defectors *u*(*D*), and down with defectors, *d*(*D*), as a function of the probability of error are plotted. As can be seen, by increasing the error probability, well close to the maximum noise level (*η* = 0.5), the densities of strategies that play soft with cooperators and those which play hard with defectors increase. While the density of those who play hard with cooperators and soft with defectors decrease. However, this increase in cooperation favoring strategies is not strong enough to compensate the loss of cooperators’ payoff in game *B* due to increasing noise in recognition. Consequently, the density of cooperators decreases when noise approaches the maximal value.

**Figure 6:**
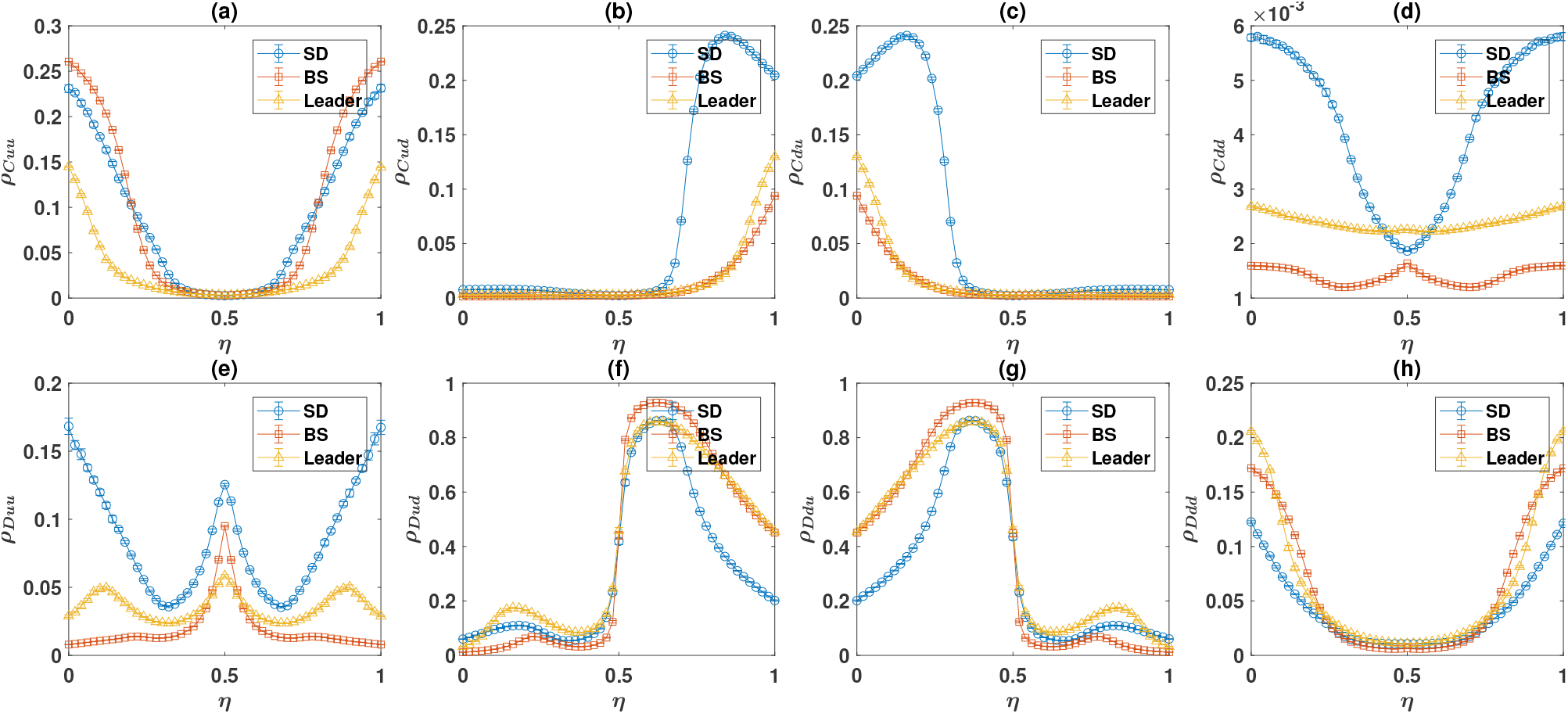
The density of different strategies in the reputation-based model with three archetypal games in a structured population. The time average density of different strategies, as a function of the probability of error in inferring the PD strategy of the opponent, *η*, are plotted. The payoff values used for the games are presented in Table. 1. Simulations are performed in a population of size 40000 individuals residing on a 200 × 200 square lattice with first nearest neighbor von Neumann connectivity and periodic boundaries. The simulations are performed for 6000 time steps, and averages and standard deviations are calculated based on the last 4000 time steps. The simulations start from random initial conditions. Here, *ν* = 0.005.

**Figure 7:**
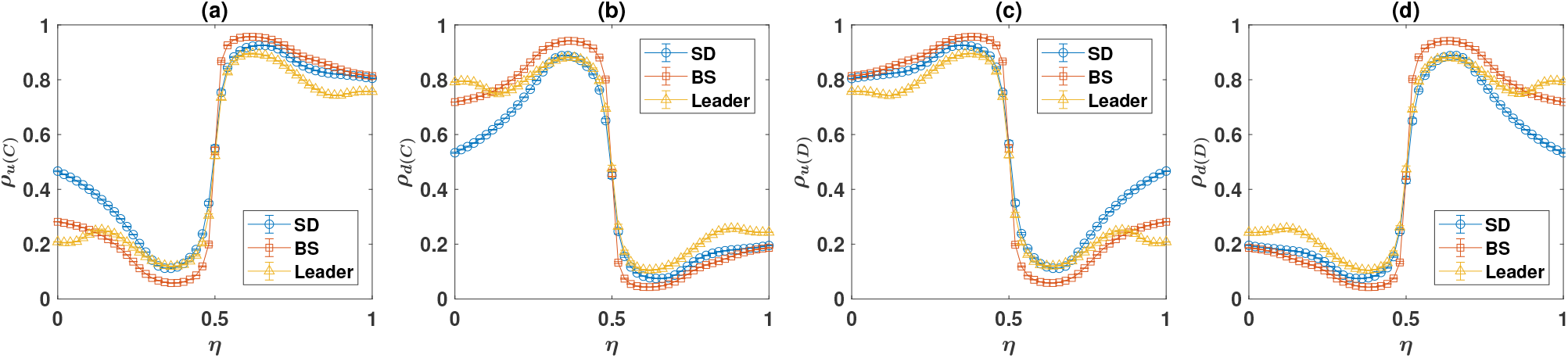
Strategic response to cooperators and defectors in game *B*, in the reputation-based model with three archetypal games in the structured population. The density of strategies who play up with cooperators *u*(*C*), down with cooperators *d*(*C*), up with defectors, *u*(*D*), and down with defectors, *d*(*D*), as a function of the probability of error in inferring the PD strategy of the opponent, *η*, are plotted. The payoff values used for the games are presented in Table. 1. Simulations are performed in a population of size 40000 individuals residing on a 200 × 200 square lattice with first nearest neighbor von Neumann connectivity and periodic boundaries. The simulations are performed for 6000 time steps, and averages and standard deviations are calculated based on the last 4000 time steps. The simulations start from random initial conditions. Here, *ν* = 0.005.

For the maximum noise level, cooperation supporting strategies experience a rapid decline (Fig. 7), and cooperation in the Prisoner’s Dilemma reaches its lowest level (Fig. 5(a)). At this point, *Dud* and *Ddu* become indiscriminate, and both dominate in the population. Interestingly, the density of cooperative strategies in game *B* reaches its maximum at this point. In the case of the Snow Drift game this value is well above the Nash equilibrium, which occurs in a mixed population. Due to the detrimental effect of network structure for the evolution of cooperation in the Snow Drift game, the density of cooperative strategies is even less than the Nash equilibrium value in a simple strategic setting in structured populations [42]. This observation shows that recognition noise can facilitate cooperation in the Snow Drift game in structured populations. We note that the mechanism behind this phenomenon seems to be rather independent from the evolution of cooperation supporting norms, as in this case, such norms do not evolve in the system.

As the probability of error increases beyond 0.5, the density of strategies that play up with defectors and down with cooperators rapidly increases (Fig. 7). Combined with the fact that in this regime, individuals are more likely to make an error in recognition of cooperators and defectors than making a correct inference, this guarantees that cooperators to be more likely to be played soft with compared to defectors. Consequently, as was the case in the mixed population, the population self-organizes into a regime where a set of cooperation supporting moral norms emerges and supports cooperation in the system.

We note that while in the mixed population, in the cooperative fixed point, *Ddu* and *Ddd* types are the only defective types which are found in large densities, in a structured population, depending on the parameter values, it can happen that all the defective strategies exist in large densities. This is due to the fact that in a structured population, domains of similar strategies are formed. While a given strategy may perform poorly globally, when surrounded by certain types, it can survive. This phenomenon, in turn, removes the bistability of the dynamics: The fate of the dynamics does not depend on the initial condition. To more closely see how this is the case, in Fig. 8(a) to 8(c) we present snapshots of the time evolution of the system, starting from a defection favoring initial condition, in which all the individuals are defectors, and a defection favoring norm prevails. That is, all the individuals are of the *Dud* type: They defect in the PD, play hard with cooperators, and soft with defectors. The time evolution of the densities of different strategies is presented in Fig. 8(d) and 8(e).

**Figure 8:**
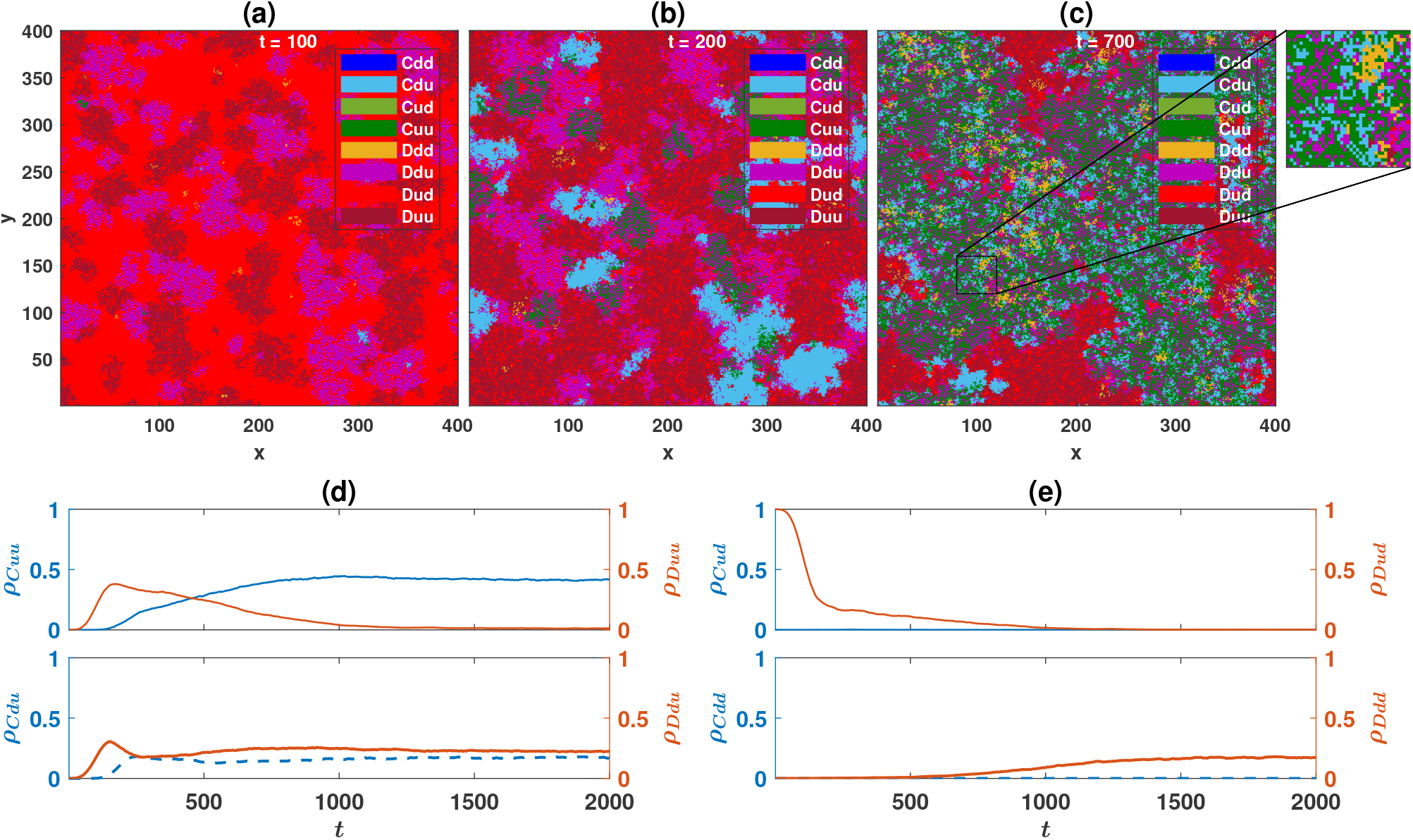
Time evolution of the system. (a) to (c): snapshots of the population during the evolution for different times, *t*, are presented. (d) and (e): The densities of different strategies as a function of time. Here, *ν* = 10^*−*4^, and game *B* is the Snow Drift game. The payoff values of the games are presented in Table. 1. The initial population is of type *Dud*. The population resides on a 400 × 400 first nearest neighbor square lattice with von Neumann connectivity and periodic boundaries.

As *Dud* individuals defer to defectors, the two defective types who play hard with defectors, *Duu* and *Ddu*, reach a higher payoff. Consequently, in the first stage of the time evolution of the system, domains of *Duu* and *Ddu* form and rapidly grow in the see of *Dud*s. However, *Dud* can coexist with *Ddu* and *Duu*. This is so because two neighboring *Ddu*s, two neighboring *Duu*s, or a *Ddu* in the neighborhood of a *Duu*, play mutually hard. On the other hand, a *Dud* by deferring to the two former strategies can perform better in their immediate neighborhood. While in a disadvantage in the see of *Dud*s, as the *Ddu* type defers to cooperators, cooperators of type *Cdu* and *Cuu*, by playing hard against *Ddu*, reach a high payoff which compensates for the cost of cooperation, and thus, *Cdu*s and *Cuu*s grow in the *Ddu* domains. This sets the second stage of the system’s time evolution, where the density of both *Cdu* and *Cuu* increases, and the density of *Ddu*, and less rapidly, *Duu* decreases. There is an important difference in the spatial patterns of the domains of *Cdu* and *Cuu* types: While *Cdu* strategies, by playing soft with each other can occupy neighboring positions, as neighboring *Cuu* types play mutually hard, they decrease the payoff of their neighbors of similar type. Consequently, *Cuu* types tend to avoid being neighbors. Instead, they form cross-like patterns where they coexist with *Ddu* type. This is crucial for the fate of the dynamics: *Cuu* and *Ddu* form a winning coalition. *Cuu* benefits *Ddu* in the PD, and *Ddu* benefits *Cuu* in the game *B*. This winning coalition can overcome the coexisting domains of *Duu* and *Dud* and invade their territories. Consequently, in the third stage of the evolution, domains of coexisting *Cuu* and *Ddu* grow at the expense of shrinking the domains of coexisting *Duu* and *Dud*. This is the stage where a cooperative norm slowly replaces a defection favoring norm. The densities of both *Duu* and *Dud* decreases, while the density of *Cuu* slowly increases in this stage.

Although, due to reaching a higher payoff in the game *B, Cuu* wins in a direct competition with *Cdu, Cuu*s benefit *Cdu*s in an indirect way. This is so because, due to the fact that *Cuu*s repel each other, *Cdu* can survive in the domains of coexisting *Ddu* and *Cuu*. Consequently, elimination of *Duu* and *Dud*, by the coalition of *Cuu* and *Ddu* benefits *Cdu* as well by increasing its territorial domain. For this reason, the density of *Cdu* slightly increases in the expansion phase of the *Cuu* and *Ddu* coalition. Finally, while *Ddd* performs poorly in the initial stages of the time evolution of the system, its density increases once a cooperative norm is established and *Dud* is removed. The reason is, in the presence of the anti-cooperative *Dud* type, *Ddd* performs poorly in indirect competition with both *Ddu* and *Duu*, who can exploit *Dud* better. However, once *Dud* is removed, *Ddd* can emerge in the system as well.

## Discussion

We have studied the evolution of strategies in a complex strategic setting, where individuals in a population play different games and base their strategy in a game on what happens in another game. By considering a situation where two interacting games exist, a Prisoner’s Dilemma followed by a second game, we have shown coupling between games can give rise to the evolution of a set of moral norms that prescribe being soft on cooperators. In this framework, as long as the second game possesses an asymmetric Nash equilibrium, it is in the individuals’ best interest to take the information about what happens in the social dilemma into account when making strategic choices in a second game. Consequently, in the course of evolution, a set of cooperation supporting moral norms emerges based on the individual’s self-interest. This appears to provide a possible explanation for the evolution of morality, in a biological population composed of self-interested individuals with simple cognitive abilities.

In our model, the cost of cooperation can be considered as a cost paid by individuals to reach a high moral status to benefit from favorable encounters in interactions that do not involve a strict social dilemma. Interestingly, the model shows, the stronger the social dilemma and the higher the cost of cooperation, a stronger set of cooperation supporting moral norms emerge, as the likelihood that cooperators receive a favorable encounter in game *B* increases with increasing the strength of the social dilemma [See Fig S.7 for a mixed population, and Fig. S.20 for a structured population]. Although this may not fully compensate the higher cost of cooperation cooperators pay in stronger dilemmas, partly alleviates cooperators’ loss of payoff and helps the evolution of cooperation when the cost of cooperation is high.

Our findings also provide insights into the evolution of indirect reciprocity. By considering a simple strategic setting, namely one in which individuals only can play a social dilemma, models of indirect reciprocity have shown that specific moral rules can support an evolutionary stable cooperative state. However, the simplicity of the strategic setting requires the moral assessment module and action module to be placed at the same level. This self-referential structure can destabilize the dynamics. To solve this problem, the theory has to appeal to higher-order and complex moral assessment rules. In addition to the lack of a natural mechanism to break the chain of higher-order rules, this requires a relatively high cognitive ability and a large amount of information about the past actions of the individuals in moral assessment [17], which appears to limit its applicability [18]. Furthermore, the dichotomy of moral assessment module and action module, commonly incorporated in many models of indirect reciprocity, can give rise to severe problems, when instead of public information, individuals have private information about the reputation of others [4,51–53]. In this case, a punishment dilemma can arise: individuals may have different beliefs about the reputation of others, and thus, disagree as to what is a justified punishment [4, 51].

In contrast, as we have shown, the introduction of strategic complexity circumvents these problems and leads to a simple dynamical mechanism for the evolution of a set of cooperation supporting moral norms. In this regard, in a strategically complex context, it is not necessary to define good and bad a priori to the dynamics of the system. This avoids the punishment dilemma when information is private. Nor it is necessary to define different moral assessment rules [10–12, 15, 16] and search for efficient ones [15–17, 19, 54]. Rather, the dynamics self-organizes into a symmetry broken cooperative phase where the symmetry between cooperation and defection breaks. A set of cooperation supporting moral norms evolves and costly cooperation emerges as a morally valuable or “good” trait due to a purely dynamical phenomenon and as a result of a symmetry-breaking phase transition.

As our analysis shows, a moral system not only promotes cooperation in a social dilemma, but it also increases cooperative behavior in a second game, a strategic setting which may be a social dilemma (as in the case of SD) or may not be a social dilemma. By considering three archetypal games, and continuous variations of the structure of the second game, we have shown this is the case for a broad range of strategic settings. In this sense, a moral system not only works to promote cooperation, but it also helps to solve coordination problems and help an efficient allocation of roles and resources. This finding seems to conform to many stylized facts about moral systems. For instance, while some moral values encourage self-sacrificing and other-regarding behavior [6, 37, 55], many other aspects of moral systems do not seem to go against individuals’ self-interest, but encourage mutually beneficial behaviors, such as mutualistic cooperation [37–41], or conflict resolution [38, 40]. Fairness, loyalty, courage, respecting others, cherishing friendship, working together, and deferring to superiors are examples of such mutualistic moral values. Based on these observations, it is suggested promoting mutually beneficial behavior can provide yet another explanation for the evolution of morality [39–41]. Importantly, in our model, this second role is what makes a moral system evolvable based solely on the individuals’ self-interest. In other words, the positive role of a moral system in bringing order and organization is beneficial at both the individual and group levels. This makes adherence to a moral system beneficial on an individual level and helps its evolution in a simple dynamical way. Interestingly, this aspect of a moral system acts like a Trojan Horse: Once established due to its organizing role, it also suppresses anti-social behavior and promotes cooperation and self-sacrifice.

Analysis of the model in a structured population confirms the robustness of our results in structured populations. Furthermore, by removing the bistability of the dynamics, population structure guarantees a cooperation supporting moral system to take over in the course of evolution even in populations initially composed of defectors with anti-cooperative norms, as long as mutations introduce new strategies in the population. In addition, noise in recognition can be beneficial for the evolution of a moral system in structured populations. Nevertheless, recognition noise also limits cooperators’ ability to benefit from stronger moral norms, and thus adversely affects cooperation. Finally, our analysis reveals in a complex strategic setting, very high levels of recognition noise facilitates the

evolution of cooperative behavior in the Snow Drift game in structured populations. This shows, in contrast to what is the case in a simple strategic setting [42], network structure can be beneficial for the evolution of cooperation in the Snow Drift game. Furthermore, this provides another case for the surprisingly beneficial role that noise may play for biological functions [43–46].

## Methods

### The replicator dynamics

The model can be solved in terms of the replicator-mutation equation, which reads as follows:

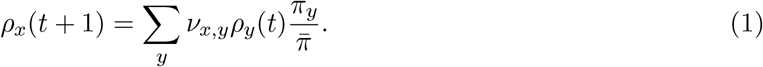

Here, *ρ*_*x*_ is the density of strategy *x, π*_*y*_ is the expected payoff of strategy *y*, 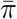 is the mean payoff, and *ν*_*x,y*_ is the mutation rate from strategy *y* to the strategy *x*. This can be written as:

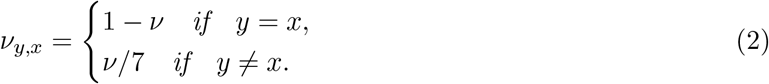

The payoff of an strategy can be written as follows. First we define:

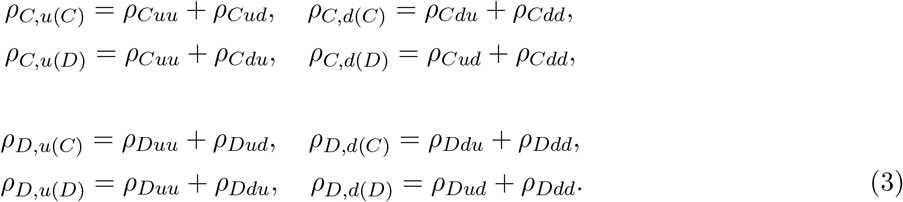

Here, the first letter in the indices shows the strategy in the PD, and *s*(*C*) (*s*(*D*)), is the strategy in the second game against a cooperator (defector). That is, for example, *ρ*_*C,u*(*C*_) is the density of those individuals who cooperate in the PD and play the up strategy with cooperators. Besides, in the following, we use *ρ*_*C*_ and *ρ*_*D*_ for the total density of those individuals who, respectively, cooperate and defect in the PD. That is, *ρ*_*C*_ = *ρ*_*Cuu*_ + *ρ*_*Cud*_ + *ρ*_*Cdu*_ + *ρ*_*Cdd*_ and *ρ*_*D*_ = *ρ*_*Duu*_ + *ρ*_*Dud*_ + *ρ*_*Ddu*_ + *ρ*_*Ddd*_.

Given these definitions, the payoffs of different strategies, in the first model can be written as follows:

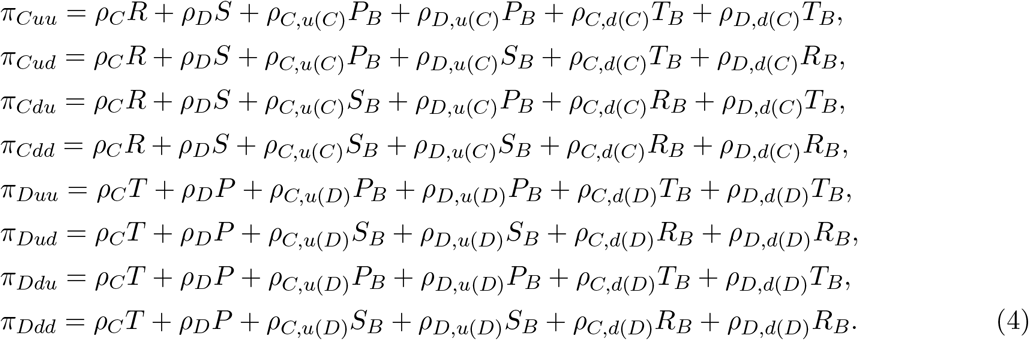

Here the first two terms in each expression are the payoffs from the PD, and the last four terms are the payoff from game *B*. The validity of these expressions can be checked by enumerating all the possible strategies that a focal individual can play with. For example, the third term in the expression for *π*_*Cuu*_ can be written by noting that a focal *Cuu* player, meets an individual of type *C, u*(*C*) with probability *ρ*_*C,u*(*C*)_. In this interaction, the focal individual plays *u* and the opponent plays *u*, leading to a payoff of *P*_*B*_ for the focal individual. Using similar arguments, it is possible to drive expressions for the payoff of different strategies in the reputation-based model. See Supplementary Information, S. 2 for details.

### Details of calculations

The base payoff values used in this study are presented in Table 1.

The density of down strategies is calculated in the simulations by directly enumerating the number of times that individuals play the strategy down in the simulation. For the solutions of the replicator dynamics these are calculated using the mean-field assumption, as *ρ*_*d*_ = *ρ*_*C*_*ρ*_*u*(*C*)_ + *ρ*_*D*_*ρ*_*u*(*D*)_. The connected correlation is calculated by assigning +1 to cooperation in the PD and strategy down in game *B*, and −1 to defection in PD and strategy up in game *B*. In the simulations, we calculate the connected correlation function, (*s*_*A*_*s*_*B*_)_*c*_ = (*s*_*A*_*s*_*B*_) − (*s*_*A*_)(*s*_*B*_), by calculating *s*_*A*_*s*_*B*_ for each individual and averaging over the population. For replicator dynamics this can be calculated using the following equation for the direct interaction model:

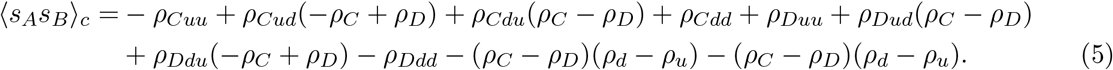

For the reputation-based model, the connected correlation function can be calculated as follows:

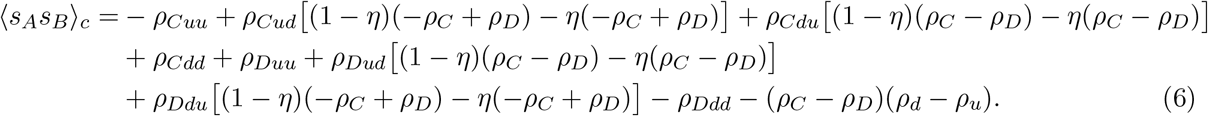

See Supplementary Information for more details.

## Acknowledgment

The author acknowledges funding from Alexander von Humboldt Foundation in the framework of the Sofja Kovalevskaja Award endowed by the German Federal Ministry of Education and Research.

## References

[1] Axelrod, R. and Hamilton, W.D., 1981. The evolution of cooperation. science, 211(4489), pp.1390–1396.

[2] Doebeli, M. and Hauert, C., 2005. Models of cooperation based on the Prisoner’s Dilemma and the Snowdrift game. Ecology letters, 8(7), pp.748–766.

[3] Nowak, M.A., 2006. Five rules for the evolution of cooperation. science, 314(5805), pp.1560–1563.

[4] Okada, I., 2020. A Review of Theoretical Studies on Indirect Reciprocity. Games, 11(3),.27.

[5] Nowak, M.A. and Sigmund, K., 2005. Evolution of indirect reciprocity. Nature, 437(7063), pp.1291–1298.

[6] Alexander, R.D., 1987. The biology of moral systems. Transaction Publishers.

[7] Nowak, M.A. and Sigmund, K., 1998. Evolution of indirect reciprocity by image scoring. Nature, 393(6685), pp.573–577.

[8] Nowak, M.A. and Sigmund, K., 1998. The dynamics of indirect reciprocity. Journal of theoretical Biology, 194(4), pp.561–574.

[9] Sugden, R., 2004. The economics of rights, co-operation and welfare (pp. 154-165). Basingstoke: Palgrave Macmillan.

[10] Leimar, O. and Hammerstein, P., 2001. Evolution of cooperation through indirect reciprocity. Proceedings of the Royal Society of London. Series B: Biological Sciences, 268(1468), pp.745–753.

[11] Panchanathan, K. and Boyd, R., 2003. A tale of two defectors: the importance of standing for evolution of indirect reciprocity. Journal of theoretical biology, 224(1), pp.115–126.

[12] Takahashi, N. and Mashima, R., 2003. The emergence of indirect reciprocity: Is the standing strategy the answer. Center for the study of cultural and ecological foundations of the mind, Hokkaido University, Japan, Working paper series, 29.

[13] Brandt, H. and Sigmund, K., 2004. The logic of reprobation: assessment and action rules for indirect reciprocation. Journal of Theoretical Biology, 231(4), pp.475–486.

[14] Kandori, M., 1992. Social norms and community enforcement. The Review of Economic Studies, 59(1), pp.63–80.

[15] Ohtsuki, H. and Iwasa, Y., 2004. How should we define goodness?reputation dynamics in indirect reciprocity. Journal of theoretical biology, 231(1), pp.107–120.

[16] Ohtsuki, H. and Iwasa, Y., 2006. The leading eight: social norms that can maintain cooperation by indirect reciprocity. Journal of theoretical biology, 239(4), pp.435–444.

[17] Santos, F.P., Santos, F.C. and Pacheco, J.M., 2018. Social norm complexity and past reputations in the evolution of cooperation. Nature, 555(7695), pp.242–245.

[18] Milinski, M., Semmann, D., Bakker, T.C. and Krambeck, H.J., 2001. Cooperation through indirect reciprocity: image scoring or standing strategy?. Proceedings of the Royal Society of London. Series B: Biological Sciences, 268(1484), pp.2495–2501.

[19] Sigmund, K., 2012. Moral assessment in indirect reciprocity. Journal of theoretical biology, 299, pp.25–30.

[20] Hilbe, C., imsa, ., Chatterjee, K. and Nowak, M.A., 2018. Evolution of cooperation in stochastic games. Nature, 559(7713), pp.246–249.

[21] Su, Q., McAvoy, A., Wang, L. and Nowak, M.A., 2019. Evolutionary dynamics with game transitions. Proceedings of the National Academy of Sciences, 116(51), pp.25398–25404.

[22] Weitz, J.S., Eksin, C., Paarporn, K., Brown, S.P. and Ratcliff, W.C., 2016. An oscillating tragedy of the commons in replicator dynamics with game-environment feedback. Proceedings of the National Academy of Sciences, 113(47), pp.E7518–E7525.

[23] Lin, Y.H. and Weitz, J.S., 2019. Spatial interactions and oscillatory tragedies of the commons. Physical review letters, 122(14), p.148102.

[24] Cressman, R., Gaunersdorfer, A. and Wen, J.F., 2000. Evolutionary and dynamic stability in symmetric evolutionary games with two independent decisions. International Game Theory Review, 2(01), pp.67–81.

[25] Chamberland, M. and Cressman, R., 2000. An example of dynamic (in) consistency in symmetric extensive form evolutionary games. Games and Economic Behavior, 30(2), pp.319–326.

[26] Hashimoto, K., 2006. Unpredictability induced by unfocused games in evolutionary game dynamics. Journal of theoretical biology, 241(3), pp.669–675.

[27] Hashimoto, K., 2014. Multigame effect in finite populations induces strategy linkage between two games. Journal of Theoretical Biology, 345, pp.70–77.

[28] Venkateswaran, V.R. and Gokhale, C.S., 2019. Evolutionary dynamics of complex multiple games. Proceedings of the Royal Society B, 286(1905), p.20190900.

[29] Donahue, K., Hauser, O.P., Nowak, M.A. and Hilbe, C., 2020. Evolving cooperation in multi-channel games. Nature communications, 11(1), pp.1–9.

[30] Wang, Z., Szolnoki, A. and Perc, M., 2014. Different perceptions of social dilemmas: Evolutionary multigames in structured populations. Physical Review E, 90(3), p.032813.

[31] Wardil, L. and da Silva, J.K., 2013. The evolution of cooperation in mixed games. Chaos, Solitons & Fractals, 56, pp.160–165.

[32] Szolnoki, A. and Perc, M., 2014. Coevolutionary success-driven multigames. EPL (Europhysics Letters), 108(2), p.28004.

[33] Amaral, M.A., Wardil, L., Perc, M. and da Silva, J.K., 2016. Evolutionary mixed games in structured populations: Cooperation and the benefits of heterogeneity. Physical Review E, 93(4), p.042304.

[34] Salahshour, M., 2019. Evolution of costly signaling and partial cooperation. Scientific reports, 9(1), pp.1–7.

[35] Salahshour, M., 2020. Coevolution of cooperation and language. Physical Review E, 102(4), p.042409.

[36] Gintis, H., Smith, E.A. and Bowles, S., 2001. Costly signaling and cooperation. Journal of theoretical biology, 213(1), pp.103–119.

[37] Gintis, H., 2013. Mutualism is only a part of human morality. Behavioral and Brain Sciences, 36(1), pp.91–91.

[38] Capraro, V. and Perc, M., 2018. Grand challenges in social physics: In pursuit of moral behavior. Frontiers in Physics, 6, p.107.

[39] Baumard, N., Andr, J.B. and Sperber, D., 2013. A mutualistic approach to morality: The evolution of fairness by partner choice. Behavioral and Brain Sciences, 36(1), pp.59–78.

[40] Curry, O.S., 2016. Morality as cooperation: A problem-centred approach. In The evolution of morality (pp. 27–51). Springer, Cham.

[41] Curry, O., Whitehouse, H. and Mullins, D., 2019. Is it good to cooperate? Testing the theory of morality-as-cooperation in 60 societies. Current Anthropology, 60(1).

[42] Hauert, C. and Doebeli, M., 2004. Spatial structure often inhibits the evolution of cooperation in the snowdrift game. Nature, 428(6983), pp.643–646.

[43] Salahshour, M., 2019. Phase Diagram and Optimal Information Use in a Collective Sensing System. Physical review letters, 123(6), p.068101.

[44] Wiesenfeld, K. and Moss, F., 1995. Stochastic resonance and the benefits of noise: from ice ages to crayfish and SQUIDs. Nature, 373(6509), pp.33–36.

[45] Zhang, H., 2018. Errors can increase cooperation in finite populations. Games and Economic Behavior, 107, pp.203–219.

[46] Ackermann, M., Stecher, B., Freed, N.E., Songhet, P., Hardt, W.D. and Doebeli, M., 2008. Self-destructive cooperation mediated by phenotypic noise. Nature, 454(7207), pp.987–990.

[47] Rapoport, A., 1967. Exploiter, Leader, Hero, and Martyr: the four archetypes of the 2 2 game. Behavioral science, 12(2), pp.81–84.

[48] Binder, K., 1987. Theory of first-order phase transitions. Reports on progress in physics, 50(7), p.783.

[49] Ohtsuki, H., Hauert, C., Lieberman, E. and Nowak, M.A., 2006. A simple rule for the evolution of cooperation on graphs and social networks. Nature, 441(7092), pp.502–505.

[50] Szab’o, G. and Fath, G., 2007. Evolutionary games on graphs. Physics reports, 446(4-6), pp.97–216.

[51] Hilbe, C., Schmid, L., Tkadlec, J., Chatterjee, K. and Nowak, M.A., 2018. Indirect reciprocity with private, noisy, and incomplete information. Proceedings of the National Academy of Sciences, 115(48), pp.12241–12246.

[52] Uchida, S., 2010. Effect of private information on indirect reciprocity. Physical Review E, 82(3), p.036111.

[53] Uchida, S. and Sasaki, T., 2013. Effect of assessment error and private information on stern-judging in indirect reciprocity. Chaos, Solitons & Fractals, 56, pp.175–180.

[54] Uchida, S., Yamamoto, H., Okada, I. and Sasaki, T., 2018. A theoretical approach to norm ecosystems: two adaptive architectures of indirect reciprocity show different paths to the evolution of cooperation. Frontiers in Physics, 6, p.14.

[55] Gintis, H., Henrich, J., Bowles, S., Boyd, R. and Fehr, E., 2008. Strong reciprocity and the roots of human morality. Social Justice Research, 21(2), pp.241–253.

